# The impact of multiple species invasion on soil and plant communities increases with invader diversity

**DOI:** 10.1101/2021.04.30.442106

**Authors:** Vujanović Dušanka, Losapio Gianalberto, Milić Stanko, Milić Dubravka

**Affiliations:** BioSense Institute, University of Novi Sad, Dr Zorana Đinđića 1, Novi Sad 21000, Serbia; Department of Biology, Stanford University, 327 Campus dr, 94305 Stanford CA, USA; Institute of Filed and Vegetable Crops, National Institute of the Republic of Serbia; Laboratory for Soil and Agroecology, Novi Sad, Serbia; University of Novi Sad, Faculty of Sciences, Department of Biology and Ecology, Trg Dositeja Obradovica 2, Novi Sad 21000, Serbia

**Keywords:** *Acer negundo*, *Amorpha fruticosa*, *Fraxinus pennsylvanica*, invasive plants, multiple invasions, soil properties

## Abstract

Despite increasing evidence indicating that invasive species are harming ecological systems and processes, impacts of multiple invasions, and the linkages between these events and changes in vegetation and soil are inadequately documented and remain poorly understood. Addressing multiple invasions would help to highlight high priority invaders and would aid in designing more effective control strategies, contributing to environmental restoration and sustainability. In this work, we tested the impact of three concurring invasive plant species, *Amorpha fruticosa, Fraxinus pennsylvanica* and *Acer negundo*, on soil conditions and native plant diversity. The research was conducted in riparian ecosystem and included the following treatments: (1) co-occurrence of the three invasive plant species, (2) occurrence of a single invasive species, and (3) control, i.e., absence of invasive species. Our findings revealed that the impact of invasive plants on soil properties and native plant diversity is magnified by their co-occurrence. Soil in mixed plots (those populated with all three invaders) contained much higher levels of nitrifying bacteria (NB), organic matter (Om), nitrogen (N), and carbon (C) as well as lower carbon to nitrogen ratio (C:N) levels, compared to single species invaded plots and control plots. Mixed plots were also characterized by reduced native plant diversity compared to single species invaded and control plots. Differences in soil conditions and native plant diversity revealed the interactive potential of invasive plants in depleting biodiversity, and thus in affecting ecological and biogeochemical processes. Our results highlight the need to study the impact of multispecies invasion and suggest that sites in riparian areas affected by co-occurring invaders, should be prioritized for ecosystem restoration.

## Introduction

One of the challenges of globalization is biotic exchange (Reaser et al., 2007). Occurring at increasing rates and volumes, the consequences of invasive species dispersed beyond their natural barriers often harm native biodiversity and impair different functions of socio-biological systems (Sala et al., 2000; Walsh et al., 2016; Qu et al., 2021). Yet, impacts of concurrent multiple invasions remain poorly understood. Elucidating individual and combined responses of invasive plants and gaining a better understanding of the impact of multispecies invasion has important ecological implications for relevant management plans and conservation of a growing number of co-invaded ecosystems.

Multispecies invasion has been identified in the vast majority of habitats (Kuebbing et al., 2013). Although multispecies invasion is potentially more detrimental to ecosystems compared to single species invasion (Simberloff and Von Holle, 1999; Inderjit et al., 2005; Pisula and Meiners, 2010), extant research on the ecological impact of invaders has primarily focused on the effects of individual invaders (Hulme et al., 2013; Kuebbing et al., 2013; D’Antonio et al., 2017; Tekiela and Barney, 2017), with studies on woody invaders being relatively scarce (Stricker et al., 2015). Co-occurrence of invasive plants can be explained by the very same introduction pathway, or by interspecific facilitation, whereby one invader facilitates the establishment and spread of another (Simberloff and Von Holle, 1999; Richardson et al., 2000; Flory and Bauer, 2014; Kuebbing and Nuñez, 2016; Zhang et al., 2020). In co-invaded ecosystems, invasive plants influence both biotic and abiotic environment, due to which their combined impact may be amplified compared to their individual effects. This joint impact on the native ecosystem results from facilitative (positive) interactions, whereby multiple species increase the magnitude of their combined response as compared to their individual responses, or competitive (negative) interactions, which occur when the combined response is weaker than a single-species response (Lortie et al., 2021). Interactions among multiple invaders can also be neutral, in which case their combined impact is missing (Kuebbing et al., 2014). Generally, interactions among co-occurring invasive plants are more commonly negative or neutral while positive interactions, although rare, are more common in woody plants and at sites with a nitrogen fixing species (Kuebbing and Nuñez, 2015). Yet, linkages among multiple invaders, native biodiversity and ecosystem properties remain overlooked.

Modification of soil properties by single or several interacting invasive plants can further support self or cross-facilitation (Vitousek, 1989; Glen and Dickman, 2005; Kuebbing and Nuñez, 2016), which in both cases hinders restoration efforts and has a subsequent cascading effect on the ecosystem. Invasive plants modify soil condition either directly by depositing leaf litter of different quality and quantity (Ehrenfeld, 2001), or indirectly by affecting the microbial communities (Kourtev et al., 2003). Although there is ample evidence that single herbaceous invaders affect soil processes and native plant communities, the impact of multiple woody invaders on soil nutrient status and native plants, especially in riparian habitats, is insufficiently documented.

The present study fills this gap in pertinent literature by examining the impact of three invasive woody species on the soil properties and native plant communities in riparian ecosystems, which due to being particularly prone to plant invasion, represent model habitats for studying ecological effects of multiple invasions (Pyšek and Prach, 1993; Planty-Tabacchi et al., 1996; Ehrenfeld and Stander, 2010). The investigation was guided by the following research questions: (1) How do the soil properties and native plant composition in plots invaded by single and three invasive plants differ from those characterizing non-invaded plots? (2) Does the impact of invasive plants increase with their richness? (3) What are the direct and indirect relationships among invasive diversity, native diversity and soil conditions?

## Materials and methods

### Study site

Our field study was conducted at a riverine wood pasture located at Krčedinska ada, which is one of the largest river islands of the Danube River basin in Serbia (covering a 2170 acre area). It is located in the northern part of Serbia, Vojvodina Province, and is a part of a larger floodplain complex and a Special Nature Reserve Koviljsko-Petrovaradinski Rit. The soil at the site is classified as Gleyic Fluvisol (IUSS Working Group WRB, 2014). The island has been used for livestock grazing for more than 100 years, which has resulted in a significant habitat and vegetation heterogeneity. The island has a history of invasion by the boxelder (*Acer negundo* L.), the green ash (*Fraxinus pennsylvanica* Marshall) and false indigo-bush (*Amorpha fruticosa* L.), which co-invade riparian areas of Eastern Europe and present major environmental management challenges. Their individual impact on ecosystems is rarely reported in the literature, while their combined impact is largely absent from available records. Their presence in the surrounding floodplain area was first recorded by Parabućski, 1972, according to whom, *F. pennsylvanica* and *A. negundo* were planted in the surrounding area, while *A. fruticosa* was probably introduced via the Danube River.

The island was surveyed during May and June of 2014 by randomly selecting 20 plots of 10 × 10 m dimensions closely located within the same flood zone, with similar soil texture (loam) and land use history, as well as the same elevation (Fig. 1), in order for the invaded plots to be as comparable as possible to uninvaded (control) plots in terms of abiotic conditions. In this way, we also minimized the probability that vegetation differed significantly within the selected plots prior to invasion. The study sample comprised of four plots populated by *A. negundo* only (AcerN), four plots with *A. fruticosa* only (AmorF), four plots with *F. pennsylvanica* only (FraxP), four plots populated by all three invasive plants (Mix), and four plots without invasive plants (Con). In all invaded plots, the abundance of invaders and the height of the individuals within a species was similar to exclude the possibility of more abundant species having an advantage compared to less abundant species, or older individuals having more time for plant-soil feedback compared to younger individuals, the trend shown by McLeod et al. (2016).

**Fig. 1.**
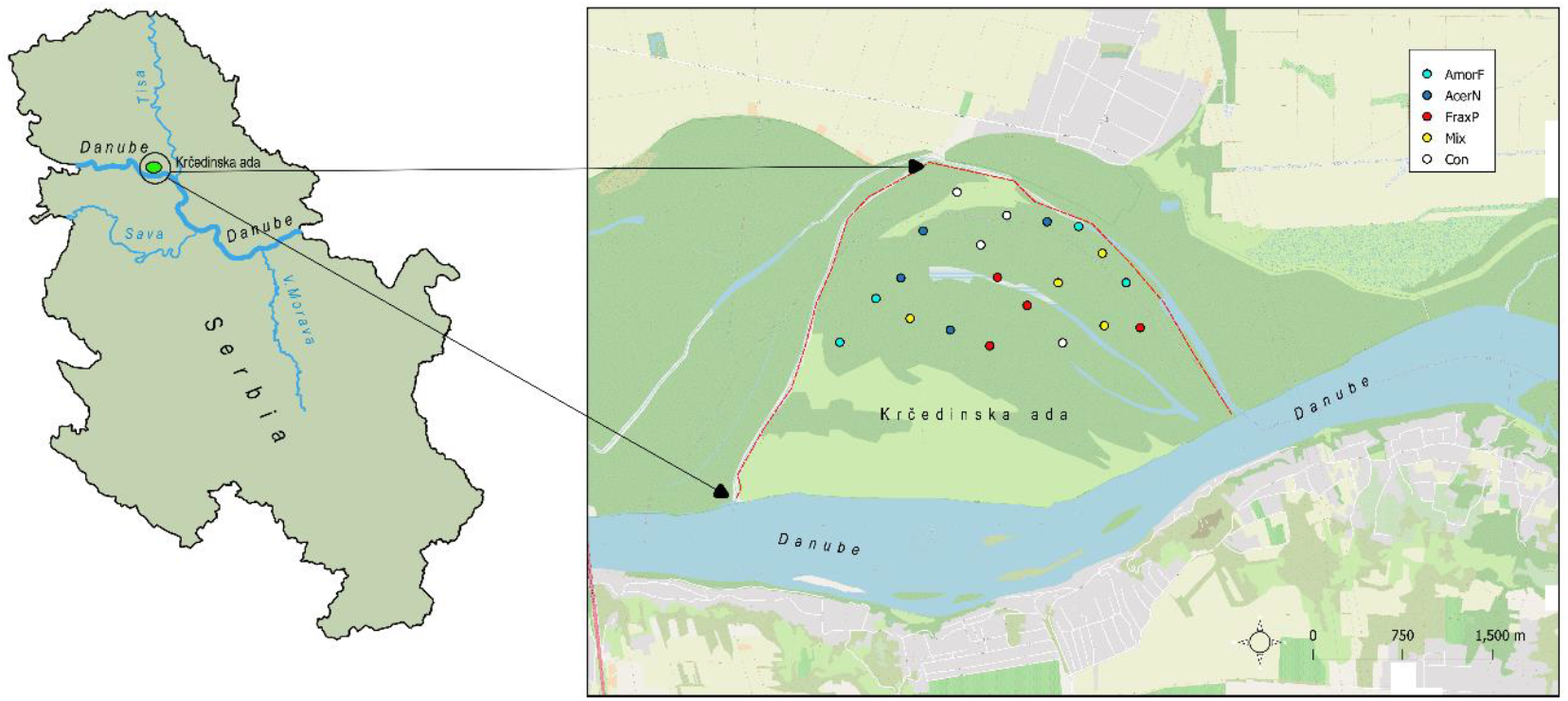
Location of the study area and the distribution of investigated plots

All vascular plant species occurring in each plot were identified to the species level, and the cover of species in each plot was visually estimated using the Braun Blanquet scale (Braun Blanquet, 1964).

### Soil analyses

Soil samples were obtained from each plot using soil probe at 0−30 cm depth, whereby one representative composite sample was formed by combining three corner samples with a 2.5 cm core diameter. Each sample was air-dried and sieved to the <2 mm particle size, in accordance with ISO 11464:2006.

All investigated soil samples were analyzed for: soil acidity (pH), calcium carbonate (CaCO_3_), organic matter (Om), plant available phosphorus AL-P_2_O_5_ (AP), plant available potassium AL-K_2_O (AK), total nitrogen (TN), carbon (C), carbon to nitrogen ratio (C:N), total sulfur (S), aluminum (Al), calcium (Ca), iron (Fe), potassium (K), magnesium (Mg), nitrifying bacteria (NB) and denitrifying bacteria (DB).

Given that all samples were of the same soil type and had uniform characteristics, to validate soil texture gradient, particle size fractions were identified in ten samples (Table S1). Particle size distribution was determined in the <2 mm fraction using the pipette method (Van Reeuwijk, 2002), revealing presence of the following size fractions: (<2 μm), silt (2−20 μm), fine sand (20−200 μm) and coarse sand (200−2,000 μm).

The soil pH value was determined in water suspension using a glass electrode in accordance with the ISO 10390:2005 methods. Calcium carbonate (CaCO_3_) content was determined in accordance with the ISO10693:1995 method for soil quality. Organic matter content was measured by the Tjurin method, while the total nitrogen and carbon content was determined via elementary analysis (CHNSO VarioEL III) in accordance with the AOAC Official Method 972.43:2006. Readily available phosphorus P (AL) and readily available potassium K (AL) in soil were determined by ammonium lactate extraction (Egner et al., 1960). Detection of available P was performed spectrophotometrically at λ = 830 nm in a UV/VIS spectrophotometer using the phosphomolybdate-blue-method (Murphy and Riley, 1962), whereas available K was determined by ammonium lactate extraction (Egner et al., 1960) using flame photometer. The total content of micro and macro elements (Mg, Fe, S, Al, and Ca) in the soil samples was analyzed after digesting the soil in concentrated HNO_3_ and H_2_O_2_ (5 HNO_3_ : 1 H_2_O_2_, and 1 : 12 solid : solution ratio) by stepwise heating up to 180 ^°^C using a Milestone Vario EL III for 55 min. Elemental concentration was determined by ICP-OES (Vista Pro-Axial, Varian) in accordance with the US EPA 200.7:2001 method. Quality control was periodically carried out with the IRMM BCR reference materials CRM-141R and CRM-142R. The recoveries were within 10% of the certified values.

For analyzing the nitrifying bacteria, the soil samples were collected aseptically from the top 30 cm layer using a hand shovel, taken to the laboratory on the same day and stored at 4 °C. Prior to the analysis, soil samples were passed through a 2mm sieve. The number of nitrifying bacteria was determined in a liquid medium by inoculating suspensions of soil dilutions in test tubes with a medium of the follwoing chemical composition: NaNO_2_ 10g; K_2_HPO_4_ 0.5g; NaCl 0.3 g; MgSO_4_ 0.5 g; MnSO_4_; Fe_2_(SO_4_)_3_; and distilled water 1000 ml. Samples were incubated for 4 days at 28 °C after which a few drops of reactive containing diphenylamine, distilled water and concentrated H_2_SO_4_ were added. The positive tubes had blue coloration. To determine the number of denitrifying bacteria soil dilution suspensions were spread directly onto nutrient agar. Gil’tai medium was used to cultivate denitrifying bacteria. After 48 hours incubation at 28 °C after which the reagents were poured over the medium and nitrate-reducing colonies were indetified by red color. The reading was converted into 1 g of soil dry weight i.e., the number of bacteria per 1 g of soil dry weight.

## Data analysis

We first tested the effects of invasive species treatments on different soil parameters and plant communities (i.e., plant alpha-diversity). We used generalized linear models (GLM) with Normal distribution for soil parameters and Poisson distribution for plant richness (seventeen separate univariate models). The former served as dependent variables while invasive species treatments were considered independent variables (categorical, with control as the reference level). Treatment significance was assesed in terms of both model fit and explained variance using Type-II ANOVA (Fox and Weisberg, 2019). Furthermore, the number of invasive species was also considered as a linear predictor, but the results remained qualitatively the same (Table S2).

Next, we tested the effects of invasive species treatments on (1) whole soil conditions, and (2) plant community composition. For this purpose, we performed sparse partial least squares discriminant analysis (sPLS–DA, Legendre and Legendre, 2012) to classify sites depending on whole soil conditions and select relevant treatments. Then, to determine the significance of treatments, we extracted the loadings of the sites along the first two axes and adopted a linear model with loadings as the dependent and treatments as the independent variable. The significance of predictors was tested both in terms of fit and 95% CI parameter estimates as well as in terms of explained variance using Type-II ANOVA (Fox and Weisberg, 2019). To visualize such a multidimensional (i.e., multivariate) dataset, results were presented as a clustered heat map and biplot.

The response of plant communities to treatments and soil conditions was subjected to cannonical correspondence analysis (CCA, Ter Braak, 1986; Legendre and Legendre, 2012), whereby plant cover data formed a community data matrix whereas invasive species treatments and significant soil properties (i.e., Om, C:N, and NB) were considered as constraining variables. Model significance of the model was tested using ANOVA-like permutation test (Legendre and Legendre, 2012).

To answer the third research question, we fitted a structural equation model (SEM, Rosseel, 2012) using the following SEM syntax: (i) regressions: native plant species diversity as a function of soil conditions and invasive diversity (i.e., number of invasive plant species), and soil conditions as a function of invasive diversity; (ii) latent variable: soil conditions as determined by the C:N ratio, Om, N, C, and NB; and (iii) correlations: C:N ratio covarying with both N and C. We used maximum likelihood estimation with robust bootstrapped SE and bootstrapped chi-squared test statistic (i.e., Satorra-Bentler correction) for model evaluation (Rosseel, 2012).

Data analysis was conducted in R 3.6.3 (R Core Team, 2020) using the ‘mixOmics’ package for sPLS– DA (Rohart et al., 2017), ‘vegan’ for CCA (Oksanen et al., 2019), and ‘lavaan’ for sem (Rosseel, 2012).

## Results

### The impact of invasion on soil and native plant diversity

Our analysis revealed significant differences in Om, N, C, C:N ratio, and NB among invasive species treatments, whereas pH, CaCO_3_, available P_2_O_5_, available K_2_O, total S, Al, Fe, Ca, K, and Mg, and denitrifying bacteria showed similar variation across treatments (Table S2).

In particular, organic matter significantly increased in *A. negundo* and *Mix* treatments as compared to control, by 33% and 53% respectively (Fig. 2a), whereas 3% and 25% increase in *A. fruticosa* and *F. pennsylvanica* was noted relative to control. Nitrogen content significantly increased in *Acer* and *Mix* treatments as compared to control, by 38% and 90% respectively (Fig. 2b), whereas 3% and 22% increase in *A. fruticosa* and *F. pennsylvanica* was noted relative to control. Carbon content significantly increased in *A. negundo, F. pennsylvanica*, and *Mix* treatments as compared to control, by 16%, 13% and 24% respectively, while 4% difference was noted between *fruticosa* and control (Fig. 2c). The C:N ratio marginally decreased (by 15%) and significantly decreased (by 34%) in *A. negundo* and *Mix* treatments as compared to control, respectively (Fig. 2d). Nitrifying bacteria showed a significant six-fold increase in *Mix* treatments compared with control (Fig. 2e). AlP_2_O_5_ significantly decreased (by 68%) in *A. negundo* while AlK_2_O marginally increased (by 52%) in *Mix* compared to control. Marginal differences were observed between *F. pennsylvanica* and control in S concentration (40% reduction). Finally, denitrifying bacteria significantly decreased (by 81%) in *A. fruticosa* relative to control. All soil samples were found to be highly calcareous (above 20%) and slightly alkaline to alkaline, as their pH ranged from 8.02 to 8.28.

**Fig. 2.**
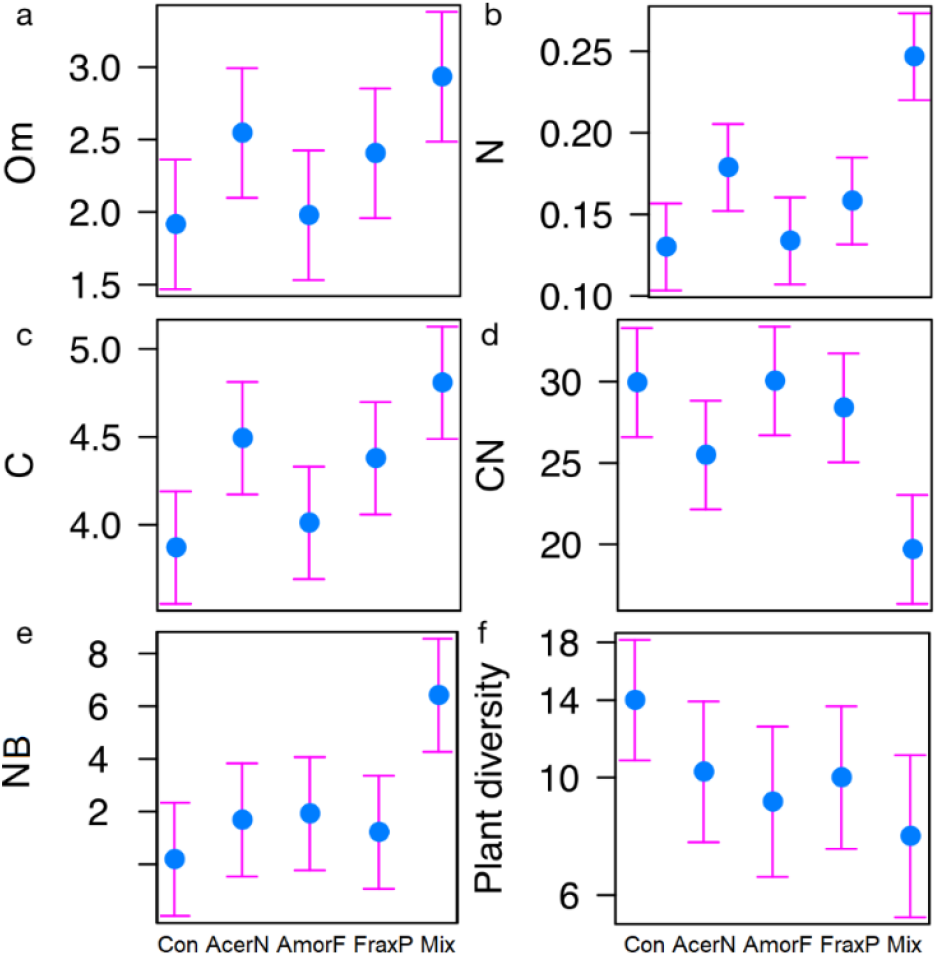
Effects of invasive species treatments (Con = control; AcerN = *A. negundo*; AmorF = *A. fruticosa*; FraxP = *F. pennsylvanica*; Mix = three-species mixture) on different soil parameters (a: organic matter; b: nitrogen; c: carbon; d: carbon to nitrogen ratio; e: nitrifying bacteria) and plant community (f: plant diversity). Estimated means and 95% CI are shown.

Looking at plant diversity, we found that invasive species treatments marginally affected the overall richness of plant species (Table S2). In particular, plant diversity significantly declined in *A. fruticosa* and *Mix* treatments as compared to control, by 17% and 22% respectively (Fig. 2f).

### Relationship among invasive species diversity, soil conditions and plant communities

The multivariate relationships among invasive species diversity, soil conditions and plant communities are shown in Fig. 3. The sPLS–DA results pertaining to the multivariate response of soil conditions to invasive species treatments, further indicate that the first and second components explained 2.9% and 14% of the variance among variables and 25.0% and 24.9% of variance among assemblages (Fig. 3a). In particular, N, C, NB and Om were positively correlated (with 0.47, 0.40, 0.38 and 0.37 correlation values, respectively), while C:N ratio was negatively correlated (a correlation of -0.43) with the first component. Moreover, Mg and CaCO_3_ were positively correlated (with 0.38 and 0.24 correlation values, respectively), while S, DB and AL-P_2_O_5_ were negatively correlated (−0.46, -0.46 and -0.34) with the second component. The distribution of assemblages along the first component reflected the actual treatments (R^2^ = 0.80, F_4,15_ = 15.00, *P* < 0.001). The results yielded by the regression analysis involving soil-condition loadings and invasive treatments indicate that control was most negatively correlated with the first axis (*β* = -1.97 ± 0.54, *P* = 0.002; -3.11 – -0.83 95% CI), and this was the only statistically significant correlation, whereas *Mix* treatment emerged as the most differential and critical one (*β* = 5.28 ± 0.76, *P* < 0.001; 3.66–6.89 95% CI). The *F. pennsylvanica* (*β* = 1.52 ± 0.76, *P* = 0.063; -0.09–3.13 95% CI) and *A. negundo* (*β* = 2.46 ± 0.76, *P* = 0.005; 0.85–4.01 95% CI) single-species treatments were also significantly and marginally associated with the first axis, respectively, while *A. fruticosa* sites exhibited an inconsistent trend (*β* = 0.58 ± 0.76, *P* = 0.455; -1.03–2.19 95% CI).

**Fig. 3.**
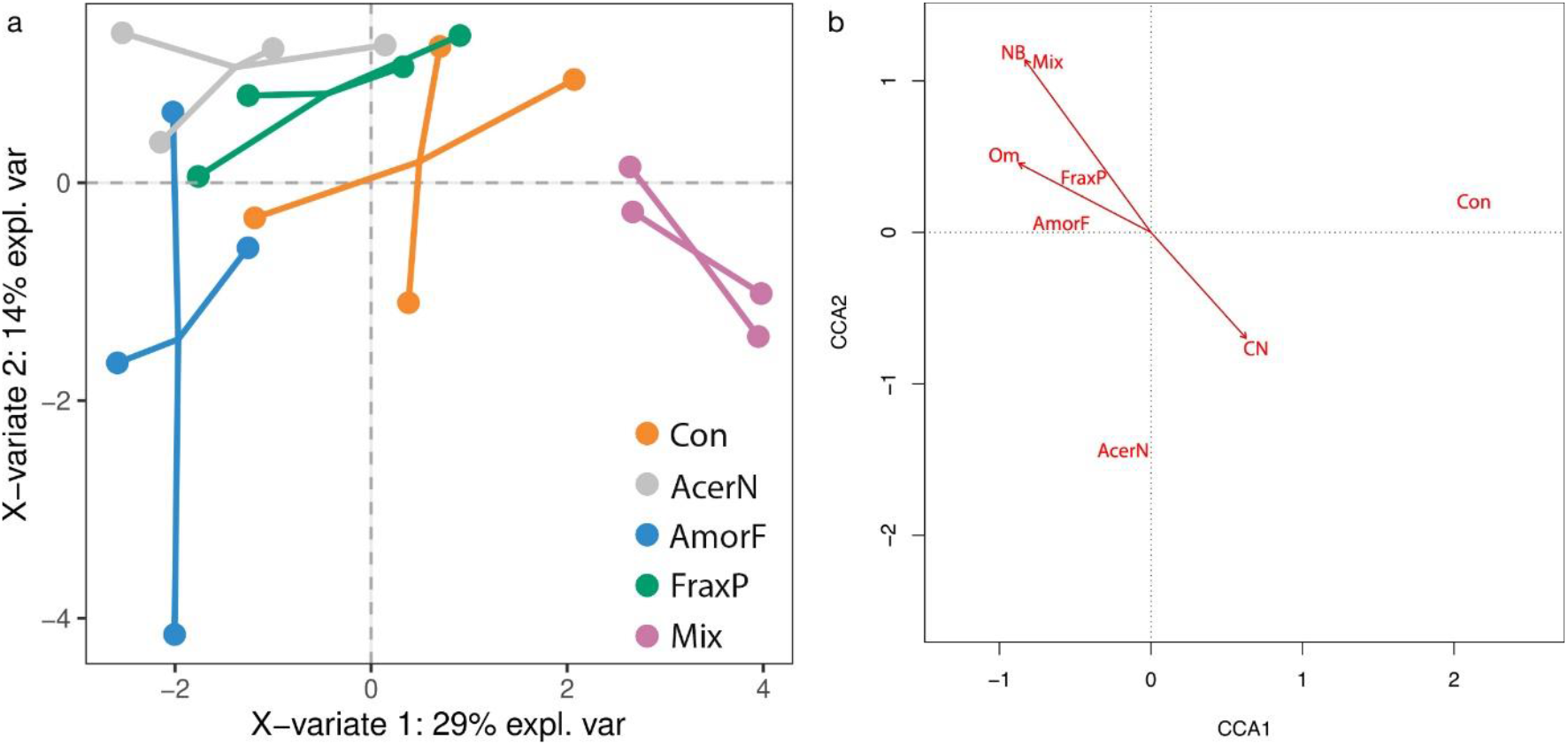
Multivariate relationships among invasive species diversity, soil conditions and plant communities. **(a)** Biplot with two main axes of variation (43% explained variance) in whole soil conditions (dots linking communities belong to the same treatment) in response to invasive treatments (orange: Control; pink: *A. negundo*; blue: *A. fruticosa*; green: *F. pennsylvanica*; gray: Mix). **(b)** Biplot showing the effects of soil conditions (arrows) on plant species distribution (omitted for clarity) across invasive treatments (red text). The first two axes explain 30% of variance.

When looking at the multivariate response of plant communities to invasive species treatment accounting for differences in soil conditions, we found that soil conditions explained 57% of variance in plant species distribution across invasive treatments (F7,12 = 1.74, *P <* 0.001; Fig. 3b). Similarly to previous results, sites were distributed following invasive treatments (*P* < 0.001) along the first axis (*P* < 0.001), with the control on one side (x_control_ = 1.89) and invasive species on the other (xacer = -0.46, xamorpha = -0.53, xfraxinus = -0.50, xmix = -0.60). Two constraining variables, Om (s = -0.49) and NB (s = -0.47) were negatively correlated, while CN (s = 0.35) was positively correlated with the first axis. The plant species most strongly associated with control were *Agropyron repens* (L.) P. Beauv., *Agrostis stolonifera* L., *Cynodon dactylon* (L.) Pers., *Diplotaxis muralis* (L.) DC., *Mentha pulegium* L., *Plantago media* L., *Plantago major* L., *Polygonum persicaria* L., *Rumex crispus* L., *Solanum nigrum* L., *Taraxacum officinale* (L.) Weber ex F.H.Wigg., *Trifolium repens* L., and *Xanthium spinosum* L. Species most strongly associated with *A. negundo* treatment were *Arctium lappa* L., *Gratiola officinalis* L., *Myosotis scorpioides* L. *Rorippa sylvestris* (L.) Besser, *Stachys palustris* L., and *Solanum dulcamara* L. Plant species most strongly associated with *Mix* treatment were *Convolvulus arvensis* L., *Galium aparine* L., *Vitis riparia subsp. longii* W.R. Prince & Prince, and *Ulmus minor* Mill. (saplings).

### Direct and indirect effect of invasion on biodiversity

Finally, we examined the direct and indirect effects of invasion on biodiversity (Fig. 4). The structural equation model (model robustness *P* = 0.149) results further indicate that invasive richness exhibited negative effects on both plant diversity (*β* = -0.96 ± 0.18, *P* < 0.001) and soil conditions (*β* = -1.01 ± 0.44, *P* = 0.022). Nevertheless, overall soil conditions had a neutral effect on plant diversity (*β* = 0.11 ± 0.43, *P* = 0.798).

**Fig. 4.**
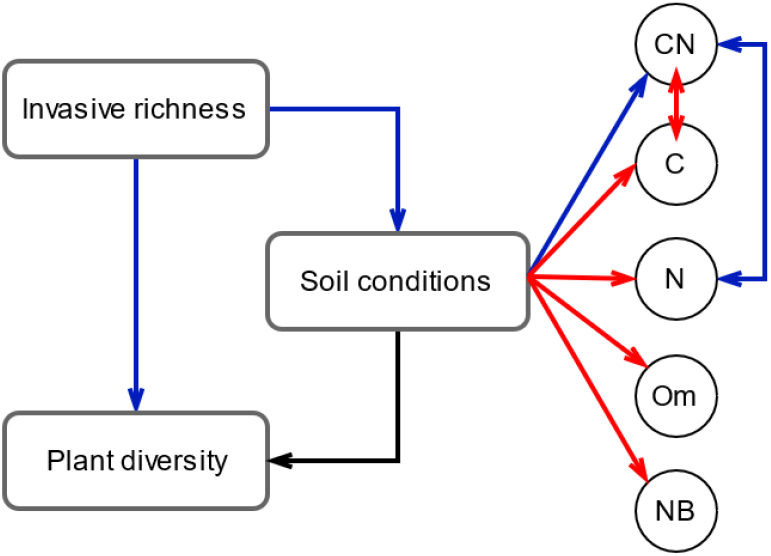
Structural equation modelling addressing the direct and indirect effects of invasive species richness on plant diversity. Blue arrows indicate negative, red arrows positive, and gray arrows neutral effects.

## Discussion

### The difference among invaded plots and uninvaded plots

The present study results indicate that differences in examined soil variables are associated with invasive plant species diversity. Notably, the impact of co-occurring invasive plants on soil properties and native plant species diversity significantly differs from the effect of single-species invasion. Our findings may be influenced by positive interactions among *A. negundo, F. pennsylvanica* and *A. fruticosa* on soil factors and native plant diversity. Longterm co-occurrence and simultaneous invasion of these three species on the studied site (Parabućski, 1972) indeed suggest their joint impact on accelerating invasion. Soils subjected to multiple invaders had significantly higher concentration of carbon, nitrogen, organic matter and nitrifying bacteria, and significantly lower carbon to nitrogen ratio, compared to non-invaded soils and those invaded by single species. These results indicate that the magnitude of the invasion impact on soil chemical properties and nitrifying bacteria content increases with multiple invader dominance.

Significant increase in the nitrifying bacteria quantity in invaded plots was detected in soils under annual grasses (Hawkes et al., 2005; McLeod et al., 2016) as well as in soils under invasive woody shrubs (Coats, 2013), compared to those under native plants. Even though we did not measure ammonium and nitrate soil concentrations, increase in the number of nitrifying bacteria in the soils under invasive plants indicates greater concentrations of these nitrogen available forms, and consequently contributes to the strengthening of competitive plant traits (Laungani and Knops, 2009; Heberling and Fridley, 2013). Feedback between plant invaders and soil microbial organisms resulting in plants securing their fitness power, is a tendency experimentally demonstrated for the nitrification process (Wolfe and Klironomos, 2005, Lee et al., 2012). Although whether increased NB content is a result of plant-microbial interactions, or plant-mediated changes in soil properties, remains to be established, our findings suggest that increased NB content in the soil is the result, rather than the driver of invasion.

The increased carbon content in the soil of *A. negundo* (16%), *F. pennsylvanica* (13%) and *Mix* (24%) plots and its slight increase in *A. fruticosa* plots (4%), support the general trend of invaders exerting direct influence on soil processes by affecting nutrient inputs through litter decomposition (Wardle et al., 2004; Liao et al., 2008; Lorenzo et al., 2010; Pyšek and Richardson, 2010; Vila et al., 2011; Si et al., 2013; Simberloff et al., 2013). Changes in soil carbon storage have been shown to be greatly enhanced by invader grouping, and could be due to a pronounced divergence in leaf and litter traits between native and invasive plants. Although we did not measure the physical and chemical leaf traits of studied species, which would enable us to predict litter decomposition rate and identify transformer species (Richardson et al., 2000), our results indicate that investigated plants belong to highly influential invaders, that are capable of modifying soil properties and nutrient cycling. Significant increase in carbon soil content at co-invaded sites may lead to imbalance in natural soil carbon stock with long-term consequences for ecosystem processes.

Nitrogen content increase compared to control was detected in *A. fruticosa* (3%), *F. pennsylvanica* (22%), *A. negundo* (38%) and *Mix* (90%) treatments. Total nitrogen content in *A. fruticosa* plots, was comparable to that in control plots, which is in accordance with the findings reported by Boscutti et al. (2020). These authors have found no significant differences in soil nitrogen content among *A. fruticosa* plots and uninvaded plots, but have found increased nitrification in soils under *A. fruticosa*, as our results have shown as well, through increased number of nitrifying bacteria in *A. fruticosa* plots. Elevated nitrogen content has been previously reported for soils under herbaceous invasive plants (Scott et al., 2001; Rodgers et al., 2008; Sanon et al., 2012) and woody shrubs (Mahla and Mlambo, 2019). The significant nitrogen increase in *Mix* plots in our study may be attributed to a higher leaf litter volume generated by *A. negundo* and *F. pennsylvanica*, which decomposes at a higher rate compared to litter produced by native plants. On the other hand, more carbohydrates for nitrogen fixing bacteria coming from high carbon input, may result in increased nitrogen soil content as well (Knops et al., 2002).

Considering that carbon and nitrogen are key macro elements, the influence of invasive plants on their respective cycles affects biogeochemical cycles of other elements. Such effect is more pronounced under prolonged invasion. Increasing soil nutrient levels, especially carbon and nitrogen, which would render the site more prone to invaders (Ehrenfeld et al., 2001), is not a trait common to all plant invaders, but is rather a characteristic of high-impact invaders, i.e., the strongest ecosystem modifiers, as shown by Jo et al. (2016).

Significantly higher increase in organic matter content in *Mix* plots, compared to single species invaded and control plots, confirms the magnified impact of these species on soil properties when co-occurring. The markedly increased organic matter content in *Mix* plots could also be attributed to a greater production of plant biomass and the resulting higher decomposition rate.

Carbon, organic matter, and nitrogen content in *Mix* plot soils may be ascribed to the positive interaction between *A. negundo* and *F. pennsylvanica*. As plant height positively correlates with the aboveground biomass and leaf mass, it is reasonable to assume that *A. fruticosa*, being a shrub, contributes less than its tree neighbors (*A. negundo* and *F. pennsylvanica*) to nutrient inputs through the leaf litter decomposition pathway.

Although native plant diversity decreased in single species-invaded plots relative to controls, the reduction was much more pronounced in *Mix* plots, demonstrating greater cumulative impact of multiple invaders.

Native plant diversity decreases due to the adverse impact of invaders on soil properties and light availability, which particularly decreased in *Mix* plots. Native species respond differently to invasion, and some are impacted more than others depending on the invader and native species traits (Stinson et al., 2007; Hejda, 2013).

Our results indicate that, when subjected to multiple invasions, native plants are displaced more rapidly compared to single species invasion. However, the findings reported by Lenda et al. (2019) counter our results, as these authors provided evidence of a much lower cumulative impact of multiple invaders on native species diversity compared to single species invasion. Such inconsistency in findings suggests that species-specific traits of invaders play a pivotal role in affecting native plant diversity. Similarly to the results of Hulme and Bremner (2006), a large number of native plants displaced by invaders in *Mix* plots are ruderal species.

### Relationship among invasive species diversity, soil conditions and plant communities

Significant differences in the soil variables, organic matter, nitrogen, carbon, carbon to nitrogen ratio, and nitrifying bacteria among invasive species treatments indicate the complexity of interspecific interactions among invasive plants.

Although soil variables characterizing *A. negundo* and *F. pennsylvanica* plots differed significantly from those measured for control plots, the greatest differences were noted in soil variables pertaining to *Mix* plots, suggesting enhanced impact of multiple invaders on soil characteristics.

Differences in soil conditions among investigated plots explained 57% of the variance in native plant distribution across investigated plots, with organic matter and nitrifying bacteria exerting the greatest influence on native plant distribution.

### Direct and indirect effect of invasion on biodiversity

Correlative relationships among invasive plant richness, plant species diversity and soil conditions, analyzed through structural equation modelling (SEM), suggest causality among invasive plant richness and soil properties. Specifically, invasive species richness exerted a direct negative effect on native plant diversity and soil properties.

According to our findings, combined impact of *A. negundo, A. fruticosa* and *F. pennsylvanica* decreased native plant diversity and negatively affected soil properties, while soil conditions had a neutral effect on plant diversity. Although negative relationship between native plant diversity and invasibility has been reported by other authors (Brown and Peet, 2003; Hejda et al., 2009; Hulme and Bremner, 2006), their investigations primarily focused on interactions between one invasive species and native vegetation.

As our findings have shown, combined impact of multiple invaders on native plants is driven by synergy between individual species-specific traits. Consequently, co-occurrence of invasive species does not always augment their impact on the ecosystem (Lenda et al., 2019).

Presence of *Ulmus minor* juveniles as well as *Convolvulus arvensis* in *Mix* plots is in line with Hejda’s (2013) observation that juveniles of tree species and species possesing a taproot tend to be more prevalent in the invaded vegetation and are least impacted by invasion.

The synergy among investigated invasive plants observed in the present study may help secure their long - term persistence, which may accelerate soil modification and native species loss, while leaving soil legacy and negatively affecting site restoration efforts. Thus, the results reported in this work can be considered when predicting harmful effects of combined invasion, and may help in mitigation of their impact on riparian sites.

Further research with additional sites and species combinations is however needed to confirm our findings and better understand invader interaction effects.

## Acknowledgments

The authors acknowledge financial support of the Ministry of Education, Science and Technological Development of the Republic of Serbia (Grant No. 451-03-9/2021-14/200358) and H2020 Project ANTARES, Grant No. 664387.

GL was supported by the Swiss National Science Foundation (P2ZHP3_187938).

We thank Csecserits Anikó, from the Institute of Ecology and Botany, Centre for Ecological Research, Alkotmány, Hungary, for constructive discussions and suggestions.

## Supporting information S1

**Table S1:** Soil texture.

| Sample | Coarse sand % | Fine sand % | Silt % | Clay % | Texture class |
| --- | --- | --- | --- | --- | --- |
| 1 | 1,94 | 44,10 | 39,72 | 14,24 | Loam |
| 2 | 2,89 | 44,27 | 38,84 | 14,00 | Loam |
| 3 | 1,07 | 68,53 | 23,20 | 7,20 | Sandy loam |
| 4 | 1,98 | 49,69 | 32,04 | 11,84 | loam |
| 5 | 2,35 | 43,77 | 40,48 | 13,40 | loam |
| 6 | 1,01 | 54,59 | 34,32 | 10,08 | loam |
| 7 | 1,48 | 58,16 | 30,28 | 10,08 | loam |
| 8 | 2,87 | 52,41 | 34,32 | 10,40 | loam |
| 9 | 2,16 | 58,24 | 30,72 | 12,88 | loam |
| 10 | 2,23 | 44,20 | 38,37 | 14,17 | loam |

## Supporting information S2

**Table S2:** Summary of regression model of soil conditions (rows) in response to invasive species treatments (columns). Model parameters are indicated as β and 95% CI estimates as lower confidence level (*lci*) and upper confidence level (*uci*). Treatments are control (*Con*), *A. negundo* (*AcerN*), *A. fruticosa* (*AmorF*), *F. pennsylvanica* (*FraxP*), and mix (*Mix*).

| | $\beta$.Con | lc.Con | uc.Con | $\beta$.AcerN | lc.AcerN | uc.AcerN | $\beta$.AmorF | lc.AmorF | uc.AmorF | $\beta$.FraxP | lc.FraxP | uc.FraxP | $\beta$.Mix | lc.Mix | uc.Mix |
| --- | --- | --- | --- | --- | --- | --- | --- | --- | --- | --- | --- | --- | --- | --- | --- |
| **pH** | 8.23 | 8.12 | 8.33 | -0.03 | -0.18 | 0.13 | -0.05 | -0.2 | 0.1 | -0.13 | -0.29 | 0.02 | -0.03 | -0.18 | 0.12 |
| **CaCO<sub>3</sub>** | 21.53 | 19.31 | 23.74 | 1.66 | -1.48 | 4.79 | 0.6 | -2.53 | 3.73 | 1.14 | -1.99 | 4.27 | -0.12 | -3.25 | 3.01 |
| **Om** | 1.92 | 1.47 | 2.36 | 0.63 | 0 | 1.26 | 0.06 | -0.57 | 0.69 | 0.49 | -0.14 | 1.12 | 1.02 | 0.39 | 1.65 |
| **ALP<sub>2</sub>O<sub>5</sub>** | 15.57 | 8.22 | 22.93 | -10.57 | -20.98 | -0.17 | -6.85 | -17.26 | 3.56 | -5.5 | -15.91 | 4.91 | -6.62 | -17.03 | 3.78 |
| **ALK<sub>2</sub>O** | 11.25 | 6.32 | 16.18 | -0.23 | -7.2 | 6.75 | -0.32 | -7.3 | 6.65 | -1.02 | -8 | 5.95 | 5.9 | -1.07 | 12.87 |
| **N** | 0.13 | 0.1 | 0.16 | 0.05 | 0.01 | 0.09 | 0 | -0.03 | 0.04 | 0.03 | -0.01 | 0.07 | 0.12 | 0.08 | 0.15 |
| **C** | 3.87 | 3.55 | 4.19 | 0.62 | 0.17 | 1.07 | 0.14 | -0.31 | 0.59 | 0.51 | 0.06 | 0.96 | 0.94 | 0.49 | 1.39 |
| **CN** | 29.94 | 26.59 | 33.29 | -4.45 | -9.19 | 0.28 | 0.1 | -4.63 | 4.83 | -1.55 | -6.28 | 3.18 | -10.26 | -14.99 | -5.53 |
| **S** | 0.06 | 0.04 | 0.08 | 0 | -0.03 | 0.02 | -0.02 | -0.05 | 0 | -0.01 | -0.04 | 0.01 | 0 | -0.02 | 0.03 |
| **Al** | 18955 | 14917.35 | 22992.65 | 702.5 | -5007.6 | 6412.6 | -860 | -6570.1 | 4850.1 | 20 | -5690.1 | 5730.1 | 1430 | -4280.1 | 7140.1 |
| **Ca** | 37685 | 35456.19 | 39913.81 | 800 | -2352.01 | 3952.01 | 232.5 | -2919.51 | 3384.51 | 117.5 | -3034.51 | 3269.51 | -535 | -3687.01 | 2617.01 |
| **Fe** | 20550 | 19106.26 | 21993.74 | 672.5 | -1369.25 | 2714.25 | 860 | -1181.75 | 2901.75 | 555 | -1486.75 | 2596.75 | 1410 | -631.75 | 3451.75 |
| **K** | 3722 | 2654.54 | 4789.46 | -118.5 | -1628.11 | 1391.11 | -514.5 | -2024.11 | 995.11 | -174.5 | -1684.11 | 1335.11 | 49.5 | -1460.11 | 1559.11 |
| **Mg** | 14880 | 14079.06 | 15680.94 | 837.5 | -295.2 | 1970.2 | 752.5 | -380.2 | 1885.2 | 802.5 | -330.2 | 1935.2 | 840 | -292.7 | 1972.7 |
| **NB** | 1844 | -19632.38 | 23320.38 | 14952.5 | -15419.69 | 45324.69 | 17342.25 | -13029.94 | 47714.44 | 10307.75 | -20064.44 | 40679.94 | 62272 | 31899.81 | 92644.19 |
| **DB** | 1627805 | 687618 | 2567992 | -572665 | -1902290.2 | 756960.2 | -1321387.5 | -2651012.7 | 8237.7 | -958002.5 | -2287627.7 | 371622.7 | -865872.25 | -2195497.45 | 463752.95 |
| **Plant diversity** | 2.64 | 2.37 | 2.89 | -0.31 | -0.72 | 0.09 | -0.44 | -0.87 | -0.03 | -0.34 | -0.75 | 0.07 | -0.59 | -1.04 | -0.16 |

## Notes

### Competing Interest Statement

The authors have declared no competing interest.

